# The Ets family transcription factor EHF suppresses senescence-associated inflammatory responses

**DOI:** 10.64898/2026.05.31.728776

**Authors:** Yukiko Fumoto, Makoto Fujikawa, Yuichiro Katayama, Theventhiran Mahandaran, Fuyuki Ishikawa, Tomoichiro Miyoshi

**Affiliations:** Laboratory for Retrotransposon Dynamics, RIKEN Center for Integrative Medical Sciences, Yokohama 230-0045, Japan; Department of Gene Mechanisms, Graduate School of Biostudies, Kyoto University, Kyoto 606-8501, Japan; Department of Pharmacology, School of Medicine, Aichi Medical University, Nagakute, Aichi 480-1195, Japan; Graduate School of Medical and Dental Sciences, Institute of Science Tokyo, Tokyo 113-8510, Japan

**Keywords:** Cellular senescence, Inflammation, Senescence-associated secretory phenotype, Ets family, NF-κB

## Abstract

Cellular senescence is a tumor-suppressive program characterized by irreversible growth arrest; however, senescent cells can also promote inflammation and alter the tumor microenvironment through the senescence-associated secretory phenotype (SASP). Although SASP induction is regulated by pathways such as p38/NF-κB/IκBζ, the mechanisms that restrain excessive or persistent SASP remain largely unknown. Here, we investigated the role of the Ets family transcription factor EHF in SASP regulation during cellular senescence. In IMR-90 human fibroblasts undergoing oncogene-induced senescence, *EHF* expression was upregulated after the onset of canonical senescence phenotypes. EHF knockdown did not substantially affect senescence establishment but increased SASP-related gene expression. Conversely, overexpression of full-length EHF suppressed SASP-related gene induction during senescence, whereas an ETS-domain-deficient EHF mutant failed to do so, suggesting that this EHF-mediated SASP suppression requires its DNA-binding domain. Furthermore, knockdown of *NFKBIZ*, which encodes IκBζ and is induced downstream of NF-κB signaling, reduced *EHF* expression during senescence; however, *NFKBIZ* overexpression increased *EHF* and SASP-related gene expression. These results link *EHF* induction to the p38/NF-κB/IκBζ inflammatory axis and support a model in which the inflammatory pathway that induces SASP also engages EHF as a negative regulator of SASP. Finally, conditioned medium from senescent cells promoted HCT116 cancer cell migration, and this activity showed a further increase after *EHF* knockdown. These findings suggest that EHF suppresses senescence-associated inflammatory responses and may function as a senomorphic effector that attenuates SASP-related inflammation without substantially affecting senescence establishment.

## Introduction

Cellular senescence was originally identified as an irreversible growth arrest caused by telomere shortening following a certain number of cell divisions in normal cells^1^. Subsequently, it was found that a similar irreversible arrest, referred to as premature senescence, was induced by diverse types of stress, including UV irradiation, DNA damage, oxidative stress, and oncogene activation^2–5^. Senescent cells typically exhibit an enlarged and flattened morphology, increased senescence-associated β-galactosidase (SA-β-gal) activity^6^, and chromatin remodeling, including formation of senescence-associated heterochromatic foci (SAHF)^7^. They also exhibit increased secretion of inflammatory cytokines, chemokines, matrix metalloproteinases (MMPs), and growth factors, collectively known as the senescence-associated secretory phenotype (SASP)^8–10^.

In addition to their autocrine effects, SASP factors secreted by senescent cells exert diverse paracrine effects on neighboring cells and tissues, including immune cell recruitment, chronic inflammation, and induction of premature senescence in adjacent cells^9,11,12^. Moreover, SASP can alter the tumor microenvironment and promote the proliferation, migration, invasion, and metastasis of cancer cells, thereby contributing to the detrimental effects exerted by senescent cells^10,13,14^. For example, IL-6, a major SASP factor, promotes tumor cell proliferation in skin cancer^15^ and breast cancer^16^, whereas MMP3, a member of the MMP family and another major component of the SASP, enhances cancer cell invasion and migration through extracellular matrix remodeling^17^. Thus, SASP is not merely a passive by-product of senescence but also a non-cell-autonomous effector program that can influence the tumor microenvironment^18^.

SASP-related gene expression is regulated by multiple signaling pathways, including NF-κB, JAK/STAT, TGF-β, and C/EBPβ, among which NF-κB plays a central role^19–22^. The NF-κB-mediated induction of SASP factors in senescent cells occurs downstream of p38 MAP kinase, a stress-responsive pathway commonly activated during senescence^23,24^. In this pathway, IκBζ, a member of the IκB family, functions as a key secondary-response transcriptional regulator controlling inflammatory SASP factors such as IL-6 and IL-8^25^. Although some negative regulators have been identified in the pathway, including miR-146a/b^26^, the molecular mechanisms that restrain excessive or persistent SASP remain largely unclear.

EHF (Ets homologous factor; also known as ESE-3), a member of the Ets family of transcription factors, is upregulated downstream of p38 activation during cellular senescence^27^. Ets family proteins bind to Ets-binding sequences, typically 5’-GGA(A/T)-3’, within regulatory regions of target genes through their Ets DNA-binding domain, thereby activating or repressing target gene transcription^28^. EHF is predominantly expressed in epithelial cells^29^ and has been implicated in differentiation and inflammatory responses^30,31^. Furthermore, overexpression of EHF induces senescence-like phenotypes in normal fibroblasts, including growth arrest and increased SA-β-gal activity, and promotes p16 transcription through Ets-binding elements in the p16 promoter^27^. Recent studies have shown that EHF induces senescence-like phenotypes in pancreatic ductal adenocarcinoma cells while suppressing SASP factor expression and influencing the tumor microenvironment^32^. In addition, NF-κB-binding sequences have been found in the promoter region of *EHF*, whose expression is activated by NF-κB in airway epithelial cells^33^ and pancreatic stellate cells^34^, suggesting that EHF may function downstream of the p38-NF-κB inflammatory axis.

Thus, accumulating evidence suggests that EHF may regulate both senescence-associated phenotypes and inflammatory responses in normal and cancer cells. However, the mechanisms governing *EHF* induction during cellular senescence, particularly its functional link to the NF-κB pathway, remain poorly understood. In this study, we therefore investigated the role of EHF in cellular senescence, focusing on its involvement in SASP regulation and its relationship to the NF-κB/IκBζ pathway.

## Materials and Methods

### Cell Culture

IMR-90 fibroblast cells (Coriell Institute for Medical Research, Camden, NJ, United States, I90-15), HEK293T (American Type Culture Collection (ATCC), Manassas, VA, United States), Platinum-A (PLAT-A; kindly provided by T. Kitamura)^35^, and HCT116 (RIKEN BRC, Ibaraki, Japan) cells were grown in Dulbecco’s Modified Eagle Medium (DMEM) (Nissui, Tokyo, Japan or Shimadzu Diagnostics, Tokyo, Japan) supplemented with 10% (volume/volume [v/v]) fetal bovine serum (FBS) (Gibco, Amarillo, TX, United States; Capricorn Scientific, Ebsdorfergrund, Germany; or MP Biomedicals, CA, United States), 0.165% (weight/volume [w/v]) NaHCO_3_ (Nacalai Tesque, Kyoto, Japan), 100 U/mL penicillin G (Sigma-Aldrich, St. Louis, MO, United States), 100 µg/mL streptomycin (Sigma-Aldrich), and 2 mM L-glutamine (Nacalai Tesque), at 37°C in a humidified atmosphere with 5% CO_2_. The absence of *Mycoplasma spp*. was confirmed using the VenorGeM Classic Mycoplasma Detection Kit (Sigma-Aldrich). Short tandem repeat (STR) genotyping was performed by BEX (Tokyo, Japan) to confirm the identity of IMR-90, HEK293T, and HCT116 cells.

### Retrovirus infection

For retrovirus production, PLAT-A packaging cells plated on a 6-cm dish were transfected with 3 μg of retroviral vector using 6 μL of PEI-MAX (40,000 MW; Polysciences, Warrington, PA, United States) in 300 μL of Opti-MEM (Thermo Fisher Scientific, Waltham, MA, United States). Viral supernatants were collected ∼48 hours after transfection and used to infect IMR-90 cells in the presence of 8 μg/mL polybrene (Sigma-Aldrich). The resulting infected cells were selected with either 2.5 μg/mL puromycin (Sigma-Aldrich) for ∼48 hours or 500 μg/mL G418 (Nacalai Tesque) for ∼96 hours.

For cellular senescence induction, IMR-90 cells transduced with *RasG12V* or *MKK6EE*-expressing retroviruses were seeded at 2.0×10^4^ cells per 6-cm dish on the day following drug selection and passaged every three days. Cells transduced with retrovirus produced from empty pMX-puro vector were used as controls for *RasG12V*- and *MKK6EE*-transduced cells. For the Raf-ER system, IMR-90 cells expressing Raf-ER, generated by transduction with pMX-puro_ΔRaf-ER vector, were treated with 100 nM 4-hydroxytamoxifen (4-OHT; Sigma-Aldrich) or ethanol (vehicle control; Nacalai Tesque) for ∼24 hours. Cells transduced with empty pMXs-neo vector were used as controls for WT-EHF- and ΔETS-EHF-expressing cells.

### shRNA transduction

Short hairpin RNAs (shRNAs) targeting *EHF*, *NFKBIZ* (protein name: IκBζ), and *LacZ* were introduced into IMR-90 cells via lentiviral infection. Each short hairpin sequence against *LacZ* (5’-AAGGCCAGACGCGAATTATTT-3’), *EHF* (5’-CGAAGACTGGTATATAAATTT-3’), or *NFKBIZ* (5’-CACTTCACATGCTGGATATTA-3’) was cloned into the lentiviral vector pCSII-U6-tet-shRNA-neo. For lentivirus production, HEK293T cells plated on a 6-cm dish were transfected with 3 μg of lentiviral vector together with 1.7 μg each of pCMV-VSV-G-RSV-Rev and pCAG-HIVgp plasmids using 6.4 μL of PEI-MAX (40,000 MW) in 300 μL of Opti-MEM. The culture supernatant containing lentivirus particles was collected, filtered through a 0.45 µm filter (Merck Millipore, Billerica, MA, United States) ∼48 hours post-transfection, and used to infect IMR-90 cells in the presence of 8 μg/mL polybrene. Infected cells were selected for ∼96 hours with 400 μg/mL G418.

### SA-β-galactosidase staining assay

IMR-90 cells were seeded at a density of 1.0×10^4^ cells/well in a 6-well plate. The following day, the cells were washed with PBS and fixed with 0.5% (w/v) glutaraldehyde (Nacalai Tesque) in PBS at room temperature for 10 min. After washing with PBS three times, the cells were incubated with Wash Buffer (1 mM MgCl_2_ [Nacalai Tesque], in PBS, pH 6.0) at room temperature for 5 min and with Staining Solution (0.56 mg/mL X-gal [5-bromo-4-chloro-3-indolyl-β-D-galactoside; Nacalai Tesque], 1 mM MgCl_2_, 5 mM K_3_Fe(CN)_6_ [Nacalai Tesque], and K_4_Fe(CN)_6_ [Nacalai Tesque] in PBS) at 37°C for 15 hours in the dark. The percentage of positively stained cells was determined by counting at least 100 cells per sample under a microscope. Images were captured using a BZ-X800 fluorescence microscope (Keyence, Osaka, Japan).

### SAHF staining

IMR-90 cells were seeded at a density of 1.0×10^4^ cells/well in a 6-well plate. The following day, the cells were washed with PBS and fixed with 4% (w/v) paraformaldehyde (Nacalai Tesque) in PBS at room temperature for 10 min. After washing three times with PBS, the cells were stained with DAPI (1 μg/mL in PBS; Nacalai Tesque) at room temperature for 10 min. The percentage of positively stained cells was determined by counting at least 100 cells per sample under a microscope. Images were captured using a BZ-X800 fluorescence microscope.

### RT-qPCR

IMR-90 cells were plated at 2.0×10^5^ cells per 6-cm dish. The following day, cells were trypsinized with 0.25% trypsin (Thermo Fisher Scientific), collected in DMEM, and washed three times with cold PBS. RNA was extracted using TRIzol (Thermo Fisher Scientific) or RNeasy Plus Mini Kit (QIAGEN, Venlo, Netherlands) according to the manufacturer’s protocol.

For TRIzol-based extraction, cell pellets were lysed in 900 μL TRIzol reagent, followed by addition of 180 μL chloroform. After vigorous shaking and incubation at room temperature for 5 min, samples were centrifuged at 12,000×g for 15 min at 4°C. The aqueous phase was transferred to a new tube, and RNA was precipitated with isopropanol (Nacalai Tesque, Kyoto, Japan). The RNA pellet was washed with 75% ethanol, air-dried, and dissolved in RNase-free water. RNA samples were then treated with TURBO DNase (Thermo Fisher Scientific) to remove genomic DNA, followed by heat inactivation. RNA was re-precipitated with ethanol and sodium acetate (Nacalai Tesque), washed, and dissolved in RNase-free water. For RNeasy-based extraction, cell pellets were lysed in Buffer RLT Plus and homogenized by vortexing for 1 min before purification, according to the manufacturer’s instructions.

Total RNA (0.5 or 1 μg) was reverse transcribed using AMV reverse transcriptase XL (TaKaRa Bio, Shiga, Japan) and oligo(dT) primers under the following conditions: 30°C for 10 min, 42°C for 30 min, and 95°C for 5 min. The resulting cDNA was subjected to quantitative PCR using Luna Universal qPCR Master Mix (New England Biolabs, Ipswich, MA, United States) on a StepOnePlus Real-Time PCR System (Thermo Fisher Scientific) or 7900HT Fast Real-Time PCR System (Thermo Fisher Scientific). For the StepOnePlus system, PCR conditions were 95°C for 15 s, followed by 40 cycles of 95°C for 15 s and 60°C for 1 min. For the 7900HT system, PCR conditions were 95°C for 1 min, followed by 40 cycles of 95°C for 15 s and 60°C for 1 min. Technical duplicates were performed for each sample analyzed. Quantification was performed by comparing the cycle threshold (Ct) values to a standard curve generated from serial dilutions of a representative sample. The primer sequences used for RT-qPCR are listed in Supplementary Table S1.

### Immunoblotting

Cell pellets were lysed in 2×SDS sample buffer (250 mM Tris [Nacalai Tesque], 4% [w/v] SDS [Nacalai Tesque], 20% [v/v] glycerol [Nacalai Tesque], and 4% [v/v] 2-mercaptoethanol [Nacalai Tesque], pH 6.8). Five micrograms of protein were separated by SDS-PAGE and transferred onto an Immobilon-FL, 0.45 μm pore, PVDF membrane (Merck Millipore) using 10 mM CAPS buffer (3-[cyclohexylamino]−1-propanesulfonic acid; Nacalai Tesque, pH 11) in a Mini Trans-Blot Electrophoretic Transfer Cell tank (Bio-Rad). After blocking in Tris-NaCl-Tween (TNT) buffer (0.1 M Tris-HCl, 150 mM NaCl, 0.1% [v/v] Tween 20, pH 7.5) with 3% (w/v) skim milk, membranes were incubated with primary antibodies at 4°C overnight. Membranes were then washed with TNT buffer, followed by incubation with the relevant secondary antibodies in TNT buffer containing 0.01% (w/v) SDS at 4°C overnight. After washing with TNT buffer, the signals were detected using an Odyssey DLx Imaging System (LI-COR, Lincoln, NE, United States) and analyzed using Empiria Studio Software version 3.2.0.186 (LI-COR). The primary antibodies used were as follows: rat monoclonal anti-ESE3 antibody (clone 5A5.5, kindly provided by A Tugores [1/100]) and mouse monoclonal anti-eIF3 p110 antibody (Santa Cruz Biotechnology, Dallas, TX, USA; sc-74507, 1/2000). The secondary antibodies used were IRDye 800CW Donkey anti-Mouse IgG (LI-COR, 926-32212, 1/10,000) and IRDye 800CW Goat anti-Rat IgG (LI-COR, 925-32219, 1/10,000). All antibodies were diluted in TNT buffer.

### Transwell migration assay

IMR-90 cells transduced with *RasG12V* or the pMX-puro empty vector were seeded at 2.0×10^4^ cells per 6-cm dish on day 9 in FBS-free DMEM. After incubation for ∼24 hours, the culture supernatants were collected, and the cell numbers were counted using a hemocytometer. The collected supernatants were normalized with FBS-free DMEM based on cell number (1.0×10^5^ cells/mL) and used as the conditioned medium. HCT116 cells were seeded onto the upper chamber of a Falcon Permeable Support for 24-well plates with an 8.0-μm pore-size PET membrane (Corning, NY, United States) at 3.0×10^3^ cells in DMEM without FBS. A total of 300 μL of conditioned medium was mixed with an equal volume of DMEM containing 20% (v/v) FBS and added to the lower chamber. Cells were incubated for ∼60 hours at 37°C in 5% CO₂. Cells remaining on the upper side of the membrane were removed, and the migrated cells on the lower side were fixed with 4% (v/v) paraformaldehyde at room temperature for 10 min, followed by staining with 1 μg/mL DAPI in PBS. Images were captured using a BZ-X800 fluorescence microscope and analyzed using the BZ-X800 Analyzer software (version 1.1.2.4, Keyence). Cell numbers were quantified using the BZ-H4M measurement software (Keyence).

### Bioinformatic analysis

Public cancer transcriptome and clinical datasets were analyzed using GEPIA (Gene Expression Profiling Interactive Analysis). Differential *EHF* expression between tumor and corresponding normal tissues was evaluated using TCGA and GTEx datasets available in GEPIA. Overall survival analysis for the selected tumors was performed using the Kaplan–Meier function in GEPIA with median *EHF* expression level using the cutoff. Statistical significance was assessed using the log-rank test.

### Statistics

For statistical analysis, comparisons between two groups were performed using an unpaired Student’s *t*-test. For comparisons among three or more groups, one-way ANOVA followed by Dunnett’s multiple comparison test was performed. For experiments involving two independent variables (e.g., time points and knockdown), two-way ANOVA followed by Tukey’s multiple-comparison test was performed. All analyses were conducted using the GraphPad Prism software (GraphPad Software, San Diego, CA, United States).

## Results

### EHF is upregulated during the course of oncogenic Ras-induced cellular senescence

Because *EHF* is markedly upregulated in senescent cells induced by various stress conditions, particularly oncogene activation^27^, oncogene-induced senescence (OIS) is a suitable model for analyzing the role of EHF in cellular senescence. We therefore examined the role of EHF in cellular senescence primarily using a senescence induction system driven by constitutively active oncogenic Ras.

When *RasG12V* was introduced into human fibroblast IMR-90 cells, cell proliferation decreased significantly from day 3 onward compared with that of control cells. By day 6, the proportion of cells positive for SA-β-gal and senescence-associated heterochromatic foci (SAHF) markedly increased (Figure 1A–D). Furthermore, the expression of SASP-related genes involved in inflammatory responses, including *IL1B*, *IL6*, *CXCL8*, and *MMP3*, also increased, peaking around day 6 after *RasG12V* introduction (Figure 1E). Thus, we confirmed that the *RasG12V*-expressing IMR-90 cells used in this study exhibited typical oncogene-induced senescence phenotypes.

**Figure 1.**
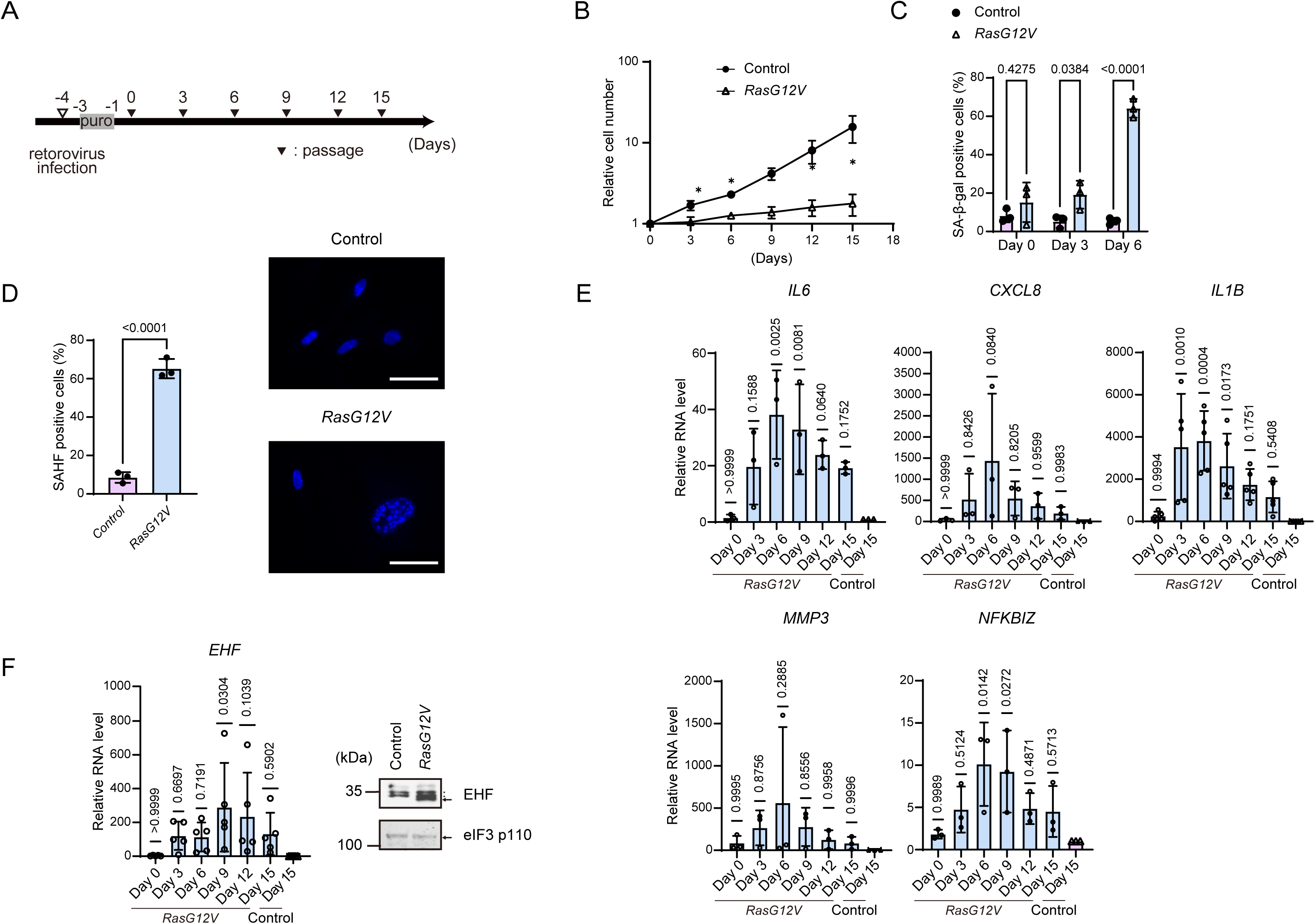
*RasG12V* induces canonical senescence phenotypes and delayed *EHF* upregulation in IMR-90 cells. A. Experimental timeline for oncogene-induced senescence (OIS) in IMR-90 cells. IMR-90 cells were infected with retrovirus containing pMX-puro_*RasG12V* or empty pMX-puro vector. After puromycin selection, the following day was defined as day 0. Cells were passaged every 3 days, and samples were collected as indicated through day 15. B. Growth curves of IMR-90 cells expressing *RasG12V* or empty vector control. Relative cell numbers were normalized to day 0. C. Percentage of SA-β-gal-positive cells on days 0, 3, and 6 after introduction of *RasG12V* or control vector. SA-β-gal-positive cells were identified by X-gal staining and counted under a microscope. D. Percentage of SAHF-positive cells on day 6 after introduction of *RasG12V* or control vector. SAHF-positive nuclei were identified by DAPI staining. E. RT-qPCR analysis of SASP-related genes in *RasG12V*-expressing cells on days 0, 3, 6, 9, 12, and 15. F. RT-qPCR analysis of *EHF* expression on days 0, 3, 6, 9, 12, and 15 after introduction of *RasG12V* or control vector, and immunoblot analysis of EHF protein on day 6. For immunoblotting, eIF3 p110 served as a loading control. For panels E and F, mRNA levels were normalized to *GAPDH* and shown relative to day-15 control cells. Statistical analyses were performed by unpaired t-test for B and D, two-way ANOVA followed by Tukey’s multiple-comparison test for C, and one-way ANOVA followed by Dunnett’s multiple-comparison test for E and F. For B, \**p* < 0.05; exact *p*-values for C–F are shown in the panels. Error bars indicate SD.

Analysis of *EHF* expression in this OIS model revealed that *EHF* mRNA levels increased over time after *RasG12V* introduction, peaking at approximately days 9–12 (Figure 1F). Western blotting also confirmed that EHF protein was already upregulated by day 6 after *RasG12V* introduction (Figure 1F). *EHF* mRNA expression became most prominent around days 9-12, later than the onset of growth arrest and increased SA-β-gal activity, which were detectable from day 3, and after SAHF formation and the peak expression of the SASP-related genes tested, which were observed around day 6. These results suggest that EHF may modulate established senescence phenotypes rather than initiating cellular senescence.

### EHF suppresses SASP-related gene expression during cellular senescence

To investigate the role of EHF in cellular senescence, we generated *EHF-knockdown* IMR-90 cells (Figure 2). At day 9 after *RasG12V* introduction, when *EHF* expression peaked in the control cells (sh*LacZ*), *EHF* mRNA levels in *EHF* knockdown cells (sh*EHF*) decreased to approximately 34% of those in control cells (sh*LacZ*). The relative decrease in *<u>EHF</u>* mRNA levels was maintained throughout the experiment (up to day 15) (Figure 2A). In contrast, sh*EHF* cells exhibited senescence phenotypes, including growth arrest and increased proportions of SA-β-gal-positive cells and SAHF-positive cells, similarly to sh*LacZ* cells (Figure 2B-C). These results suggest that EHF is not required for establishing cellular senescence.

**Figure 2.**
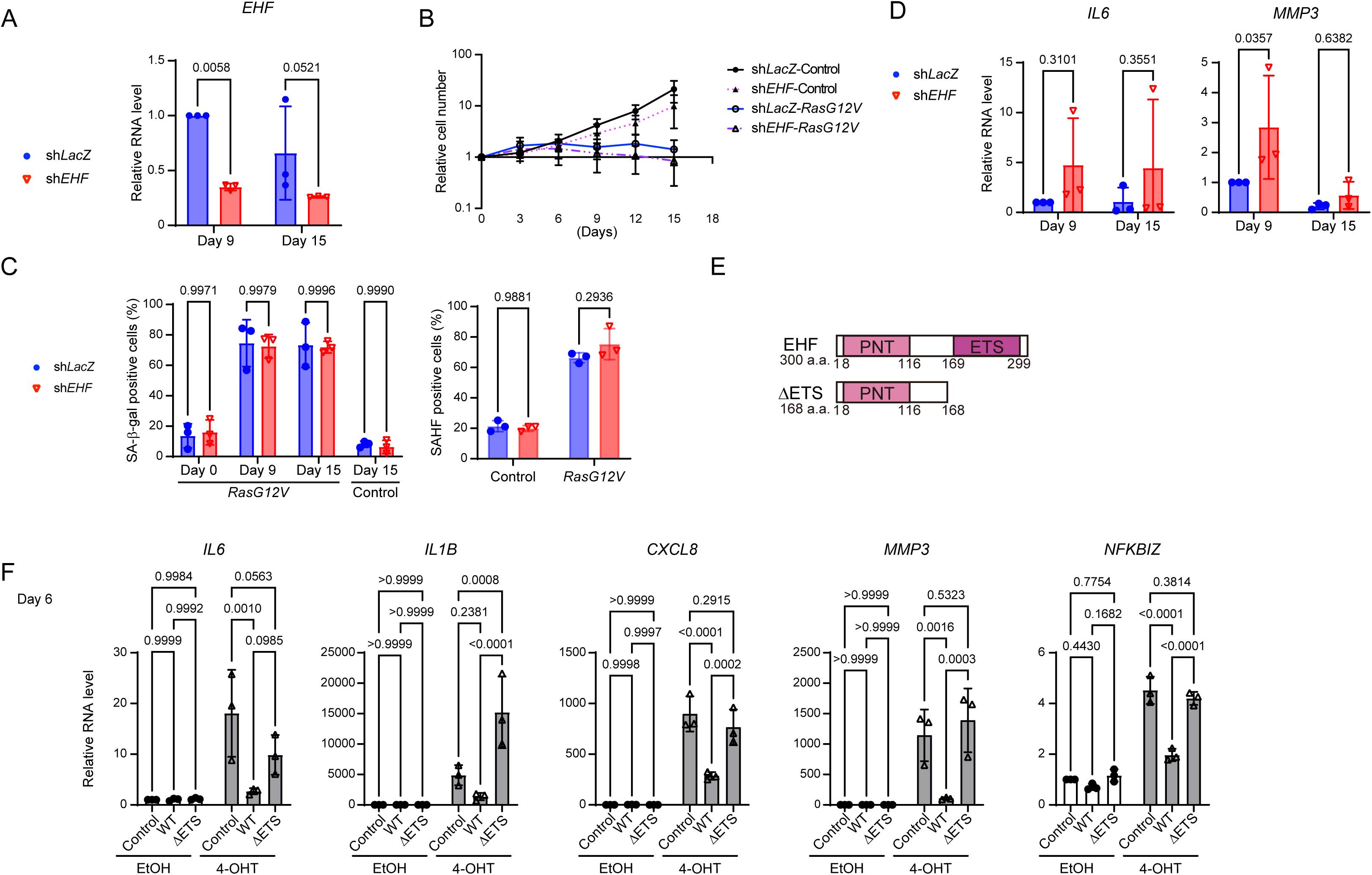
EHF suppresses senescence-associated induction of SASP-related genes. A. RT-qPCR analysis of *EHF* knockdown efficiency in sh*LacZ*- and sh*EHF*-expressing IMR-90 cells after introduction of *RasG12V* or control vector. *EHF* expression levels were measured on days 9 and 15, and shown relative to sh*LacZ-*expressing cells on day 9. B. Growth curves of sh*LacZ-* and sh*EHF*-expressing cells after introduction of *RasG12V* or control vector. Relative cell numbers were normalized to day 0. C. Percentages of SA-β-gal-positive cells and SAHF-positive cells in sh*LacZ-* and sh*EHF*-expressing cells upon *RasG12V*-induced senescence. SA-β-gal staining was performed on the indicated days, and SAHF formation was determined by DAPI staining on day 6. D. RT-qPCR analysis of *IL6* and *MMP3* in sh*LacZ-* and sh*EHF*-expressing cells after senescence induction. Cells were collected on days 9 and 15. mRNA levels were shown relative to sh*LacZ*-expressing cells on day 9. E. Schematic of full-length EHF (WT) and the ETS domain-deleted mutant ΔETS. Full-length EHF contains the N-terminal PNT domain and the C-terminal ETS DNA-binding domain, but ΔETS lacks the ETS domain. F. RT-qPCR analysis of SASP-related genes and *NFKBIZ* expression in vector-control, WT-EHF, and ΔETS-EHF-expressing cells upon Raf-ER-induced senescence. Cells were treated with 100 nM 4-OHT or EtOH for 24 hours and cultured for 5 additional days before RNA extraction. mRNA levels were shown relative to the corresponding EtOH-treated control. Senescence-associated gene induction in the Raf-ER senescence system is shown in Supplementary Figure S1. For panels A, D, and F, mRNA levels were normalized to *GAPDH*. Statistical analyses for A, C, D, and F were performed by two-way ANOVA followed by Tukey’s multiple-comparison test. Exact *p*-values are shown in the panels. Error bars indicate SD.

Previous studies have shown that EHF regulates inflammatory responses in various tissues and cell types^29–31,36^. Given that *EHF* expression peaks slightly later than the establishment of senescence phenotypes and the timing of peak expression of SASP-related genes in this model (Figure 1E–F), we hypothesized that EHF modulates SASP gene expression during cellular senescence in normal human fibroblasts. Indeed, *MMP3* expression was markedly increased in sh*EHF* cells compared to that in control sh*LacZ* cells in the RasG12V-induced senescence experiments (Figure 2D). Furthermore, although not statistically significant, *IL6* expression was higher in sh*EHF* cells than in sh*LacZ* cells. These results suggest that EHF suppresses the upregulation of SASP-related genes during cellular senescence.

To corroborate this possibility, we examined SASP gene expression in *EHF-*overexpressing cells. Because *EHF* overexpression has been shown to delay cell proliferation^27^, it was possible that the procedure compromised the efficiency of retrovirus-mediated *RasG12V* introduction. Therefore, we instead used a Raf-ER-inducible senescence system^23,37^. In this system, 4-OHT-treated cells were compared with their corresponding EtOH-treated controls for each construct. On day 6 after 4-OHT-mediated Raf-ER activation in IMR-90 cells, *CDKN1A* and *MMP3* were significantly upregulated, and *IL6*, *CXCL8*, and *EHF* also showed upward trends, indicating that this system induced senescence-associated gene expression in IMR-90 cells, similarly to the *RasG12V*-induced condition (Supplementary Figure S1). In cells overexpressing full-length EHF, *EHF* mRNA levels remained higher than those in vector control cells (pMXs-neo), both before and after senescence induction (Supplementary Figure S1C). Under these conditions, the induction of SASP-related genes (*IL1B*, *IL6*, *CXCL8*, and *MMP3*) was suppressed on day 6 after senescence induction (Figure 2F). In contrast, this suppression was not observed in IMR-90 cells expressing an EHF mutant lacking the ETS domain required for DNA binding. Taken together, these results indicate that EHF suppresses the senescence-associated induction of SASP-related genes, and that this suppression requires the DNA-binding domain of EHF (Figure 2F).

### The p38/NF-κB/IκBζ axis contributes to the induction of SASP-related genes and *EHF*

In cellular senescence, p38 MAP kinase has been reported to upregulate *EHF* expression^27^. However, EHF has not been reported as a direct substrate of MAP kinases^38^, and the underlying mechanism of p38-mediated EHF activation remains unclear. Although multiple signaling pathways contribute to the activation of SASP-related genes, the NF-κB pathway has been widely recognized as the major axis of senescence^22,24^. In this context, IκBζ (encoded by *NFKBIZ*), a nuclear IκB family member downstream of NF-κB, contributes to the transcriptional activation of SASP-related genes, such as *IL6* and *CXCL8*^25^. Activated NF-κB induces *NFKBIZ* expression, and the resulting increase in IκBζ subsequently promotes the expression of SASP-related genes. Thus, this pathway represents a secondary transcriptional response downstream of NF-κB activation. Consistent with this idea, *NFKBIZ* expression increased during senescence induced by *RasG12V* or Raf-ER at day 6 (Figures 1E and Supplementary Figure S1B). Furthermore, in cells expressing MKK6EE, a constitutively active mutant of MKK6 that functions upstream of p38, the expression levels of *NFKBIZ* and *EHF* were also elevated, together with those of *IL1B*, *IL6*, *CXCL8*, *MMP3*, and *CDKN1A* (Figure 3A and Supplementary Figure S2B)^23,24,27^. In contrast, no significant changes were observed in the mRNA levels of *NFKB1* (p50) and *RELA* (RelA), which encode the NF-κB subunits. These observations suggest that activation of the p38 pathway induces *EHF* expression through a secondary NF-κB/IκBζ-dependent transcriptional cascade, rather than by increasing the expression of NF-κB subunits.

**Figure 3.**
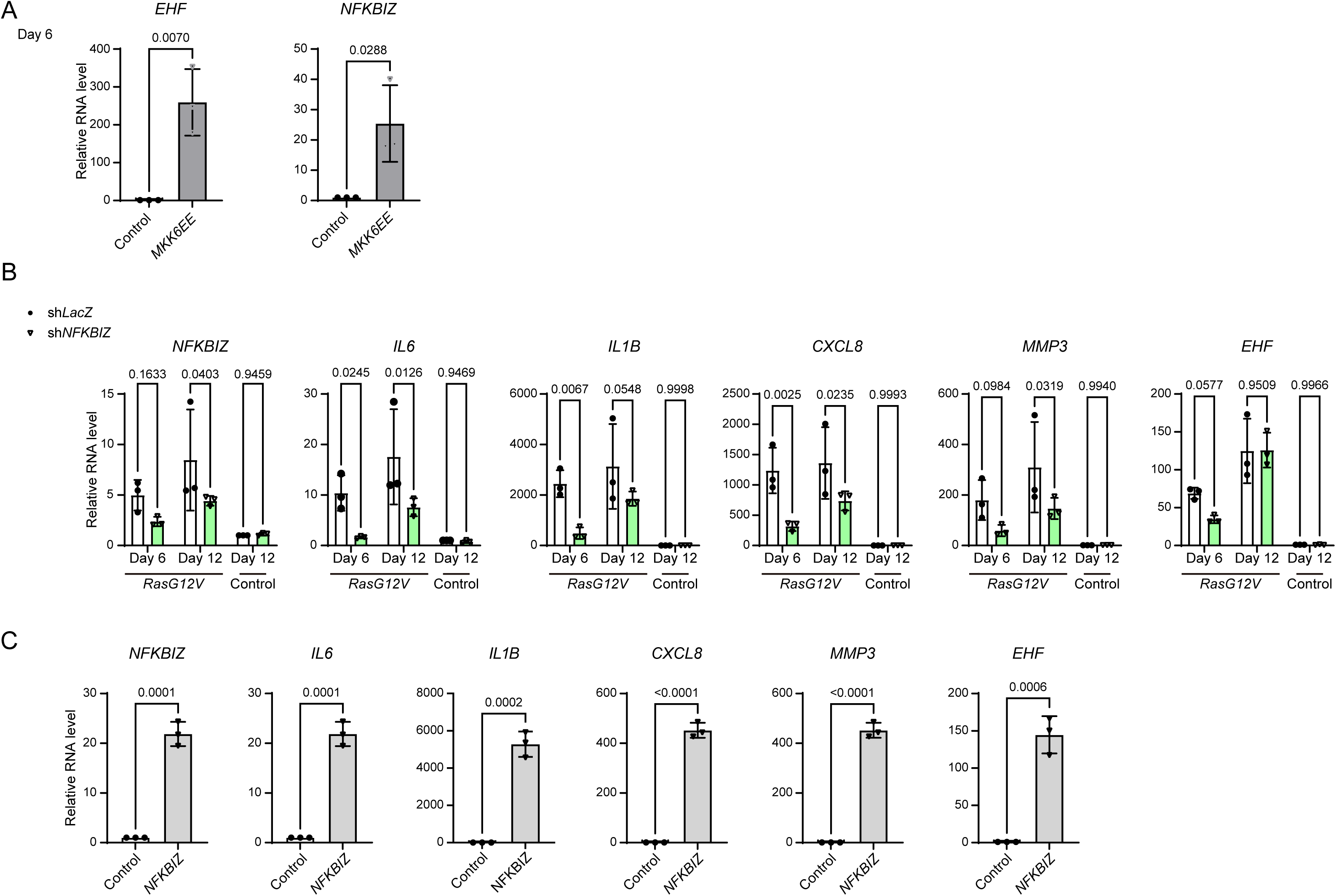
The p38/NF-κB/IκBζ pathway is involved in inducing SASP-related genes and *EHF* A. RT-qPCR analysis of *EHF* and *NFKBIZ* after expression of constitutively active MKK6EE in IMR-90 cells. Cells transduced with the empty pMX-puro vector served as controls. Cells were collected on day 6 after transduction. mRNA levels were shown relative to control cells. B. RT-qPCR analysis of *NFKBIZ*, *EHF*, and SASP-related genes in sh*LacZ-* and sh*NFKBIZ*-expressing cells upon *RasG12V*-induced senescence. Cells were collected on the indicated days after induction. mRNA levels were shown relative to sh*LacZ*-expressing cells transduced with the control vector. C. RT-qPCR analysis of *NFKBIZ*, *EHF*, and SASP-related genes in proliferating IMR-90 cells overexpressing *NFKBIZ* (IκBζ). mRNA levels were shown relative to control cells. For A–C, mRNA levels were normalized to *GAPDH*. Statistical analyses were performed by an unpaired t-test for A and C and by two-way ANOVA followed by Tukey’s multiple-comparison test for B. Exact *p*-values are shown in the panels. Error bars indicate SD.

To test this possibility, we generated *NFKBIZ* knockdown IMR-90 cells and performed RT-qPCR analysis of selected senescence-related genes and *EHF* during *RasG12V*-induced senescence (Figure 3B). In these cells, the expression levels of SASP-related genes (*IL1B*, *IL6*, *CXCL8*, and *MMP3*) decreased compared with those in sh*LacZ* control cells after *RasG12V* introduction, consistent with a previous report^25^. Furthermore, *EHF* expression was reduced in sh*NFKBIZ* cells compared with that in sh*LacZ* cells on day 6 after senescence induction. Conversely, when *NFKBIZ* was overexpressed in proliferating IMR-90 cells, both *EHF* and SASP-related genes were significantly upregulated (Figure 3C). Moreover, in senescent cells overexpressing *EHF*, *NFKBIZ* mRNA levels were reduced (Figure 2F). Taken together, these results suggest that IκBζ contributes not only to the transcriptional activation of SASP-related genes but also to *EHF* upregulation. We propose a model in which, during cellular senescence, the upstream stress-responsive pathway involved in SASP induction may subsequently engage *EHF*-mediated suppression.

### Conditioned medium from *EHF* knockdown senescent cells tends to enhance migration of neighboring cancer cells

SASP factors released from senescent cells act in a paracrine manner on neighboring cells and possibly enhance malignant phenotypes, including cancer cell migration and invasion^10,14^. Therefore, we examined the effect of IMR-90-derived conditioned medium on the migration of the human colon cancer cell line HCT116 (Figure 4A)^39^. The conditioned medium from *RasG12V*-induced senescent cells increased the number of HCT116 cells migrating through the transwell compared with the conditioned medium from control cells (Figure 4B). In the donor IMR-90 cells used for conditioned medium collection, mRNA levels of SASP-related genes (*IL1B*, *IL6*, *CXCL8*, and *MMP3*) were increased at the time of harvest (Supplementary Figure S3A). These results indicate that the conditioned medium from senescent cells with upregulated SASP gene expression promotes the migration of neighboring cancer cells, as reported previously^13,17^.

**Figure 4.**
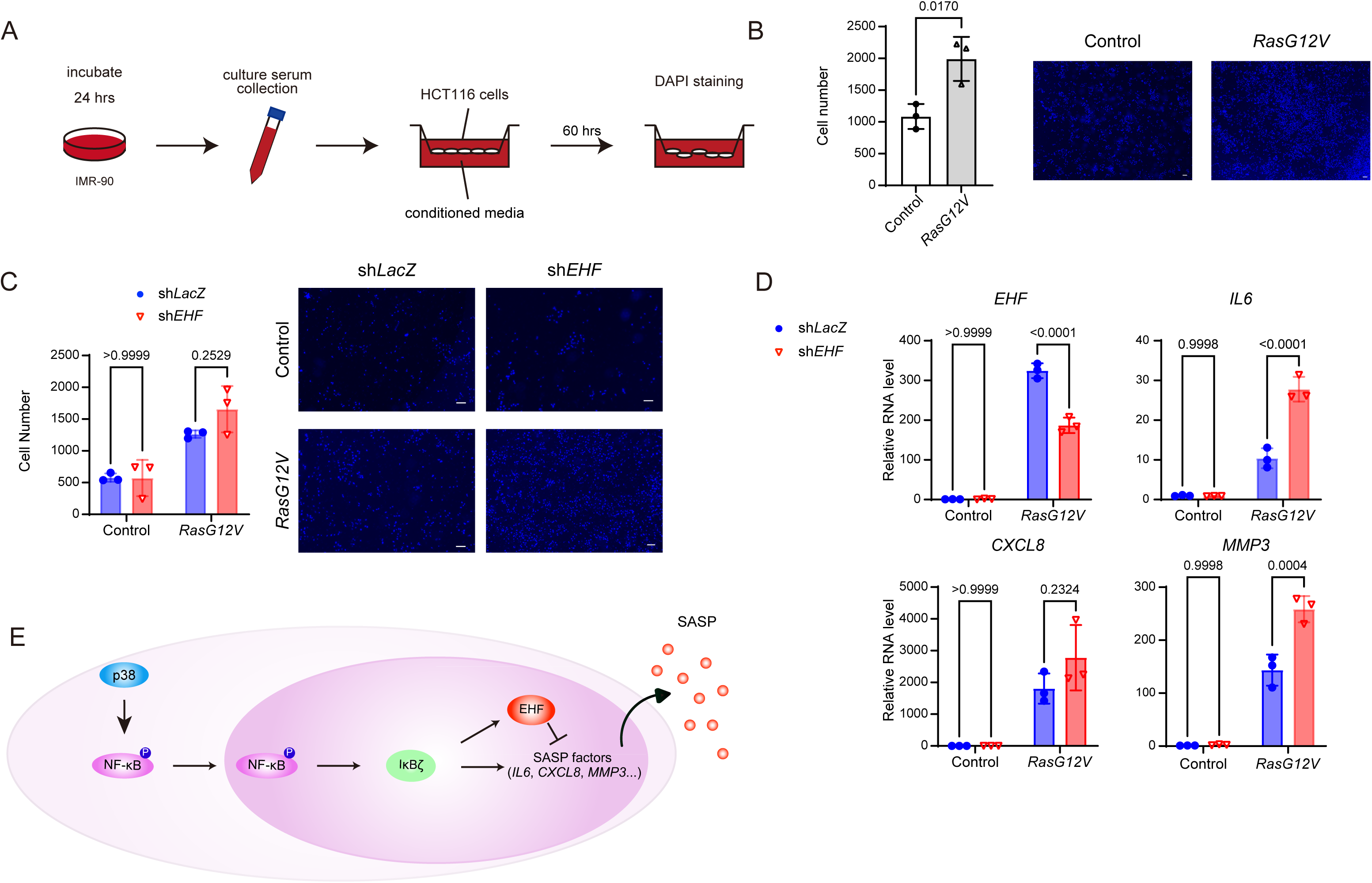
EHF attenuates the pro-migratory activity of conditioned medium from senescent cells. A. Schematic of the transwell migration assay. Conditioned medium was collected from IMR-90 donor cells after 24 hours of culture in serum-free DMEM and adjusted to 1×10^5^ donor cells/mL based on donor-cell number. HCT116 cells were seeded in the upper chamber of an 8.0-μm pore transwell insert, and migration toward conditioned medium in the lower chamber was assessed after 60 hours. Migrated cells on the lower surface were fixed, stained with DAPI, and counted. B. Migration of HCT116 cells in conditioned medium prepared from IMR-90 cells expressing *RasG12V* or empty vector. Conditioned medium was collected on day 9 after *RasG12V* or a control vector was introduced. Migrated HCT116 cell numbers are shown on the left, and representative images of migrated cells are shown on the right. Scale bar, 100 μm. C. Migration of HCT116 cells in conditioned medium prepared from sh*LacZ-* or sh*EHF*-expressing IMR-90 cells transduced with *RasG12V* or control vector. Migrated HCT116 cell numbers are shown on the left, and representative images of migrated cells are shown on the right. D. RT-qPCR analysis of *EHF*, *IL6*, *CXCL8*, and *MMP3* in donor IMR-90 cells at the time of conditioned-medium collection for the experiment shown in C. mRNA levels were normalized to *GAPDH* and shown relative to control-vector-transduced sh*LacZ-*expressing cells. Statistical analyses were performed by an unpaired t-test for B and by two-way ANOVA followed by Tukey’s multiple-comparison test for C and D. Exact *p*-values are shown in the panels. Error bars indicate SD. E. Working model for EHF-mediated suppression of SASP-related gene expression during cellular senescence. During cellular senescence, p38 MAPK activation promotes NF-κB signaling, leading to induction of *NFKBIZ* (IκBζ). Upregulated IκBζ further promotes expression of SASP-related genes, including *IL6*, *CXCL8*, and *MMP3*, as well as *EHF*. In turn, EHF suppresses SASP-related gene expression at the transcriptional level.

Next, we examined the conditioned medium prepared from control knockdown (sh*LacZ*) and *EHF*-knockdown (sh*EHF*) cells after *RasG12V*-induced senescence. The migration of HCT116 cells increased in the conditioned medium from *RasG12V*-induced sh*LacZ* cells and appeared to further increase, although this increase was not statistically significant, in the conditioned medium from sh*EHF* cells (Figure 4C). In the *RasG12V*-induced sh*EHF* donor cells, *EHF* mRNA levels were reduced, *IL6* expression was significantly increased, and *CXCL8* and *MMP3* expression showed upward trends (Figure 4D). These results suggest that the upregulation of SASP-related genes following *EHF* knockdown may contribute to the migratory response of neighboring cancer cells. However, in one of the three independent experiments, the conditioned medium from sh*EHF* cells did not clearly enhance HCT116 migration compared with that from control cells (Supplementary Figure S3B). In this replicate, we found that while *IL6* and *MMP3* expression remained upregulated, *CXCL8* expression did not increase (Supplementary Figure S3C). It is possible that the migratory response depends not only on the overall increase in SASP gene expression, but also on the activation of a specific combination of SASP-related factors, potentially including *CXCL8*. Future studies are required to examine this possibility.

## Discussion

This study demonstrated that *EHF* is upregulated late during oncogene-induced senescence and suppresses SASP-related gene expression after senescence has been established. In *RasG12V*-expressing IMR-90 cells, *EHF* expression peaked after canonical senescence phenotypes, such as growth arrest, increased SA-β-gal activity, and SAHF formation, had become evident (Figure 1). Furthermore, *EHF* knockdown increased SASP-related gene expression without markedly affecting senescence establishment, whereas *EHF* overexpression suppressed it in an ETS-domain-dependent manner (Figure 2). These results suggest that EHF functions as a transcriptional regulator that attenuates the inflammatory phenotype in oncogene-induced senescence rather than driving senescence entry (Figure 4E).

We previously showed that *EHF* overexpression induces premature senescence, as evidenced by increased p16 expression and delayed proliferation^27^. In contrast, *EHF* knockdown did not impair the establishment of typical senescent phenotypes (Figure 2). This discrepancy suggests that senescence entry is controlled by multiple transcriptional regulators, including other Ets family transcription factors such as Ets1/2. Because Ets family members recognize similar DNA-binding motifs, their target gene binding and transcriptional functions may partially overlap^40,41^. Thus, under proliferative conditions, *EHF* overexpression may act dominantly on senescence-associated genes such as p16, whereas, in oncogene-induced senescence, where multiple Ets factors are activated^42,43^, EHF appears to function mainly in suppressing SASP-related gene expression after senescence is established.

Our data further support the involvement of the p38/NF-κB/IκBζ axis in *EHF* induction. Activation of p38 by MKK6EE led to increased expression of *IL1B*, *IL6*, *CXCL8*, *MMP3*, *CDKN1A*, *NFKBIZ,* and *EHF*. In contrast, *NFKBIZ* knockdown reduced expression of SASP-related genes and *EHF* during *RasG12V*-induced senescence, whereas *NFKBIZ* overexpression increased both SASP-related gene and *EHF* expression in proliferating IMR-90 cells (Figure 3). Thus, IκBζ contributes not only to SASP-related gene expression but also to *EHF* upregulation, suggesting that *EHF* acts downstream of a secondary NF-κB/IκBζ-dependent transcriptional response.

In the NF-κB pathway, negative feedback regulators, such as miR-146a/b, are induced by SASP-inducing signaling and suppress excessive inflammatory responses^26^. Similarly, in our system, *EHF* expression kinetics suggest that *EHF* is not a trigger of SASP but a delayed suppressor that limits its persistence. However, this pathway is unlikely to be unidirectional, because *NFKBIZ* expression was reduced in cells overexpressing *EHF* in addition to SASP-related genes (Figure 2). This suggests that EHF not only suppresses SASP-related genes but may also influence the upstream transcriptional network. Thus, the p38/NF-κB/IκBζ/EHF axis may function as a regulatory network in which both SASP-facilitating and SASP-restricting pathways operate in parallel with a temporal delay. Because this study did not directly analyze genomic binding of EHF, whether EHF directly represses individual SASP-related genes or *NFKBIZ* remains to be determined. The delayed kinetics of *EHF* induction are consistent with a negative regulatory module; however, direct evidence that EHF forms a time-dependent feedback circuit with the NF-κB/IκBζ pathway will require further analyses.

Previous studies also suggest that EHF regulates gene expression in a context-dependent manner^38^. In PDAC cells, EHF promotes a senescence-like phenotype while suppressing SASP gene expression^32^, consistent with our findings. In contrast, in pancreatic stellate cells, EHF upregulates IL-6 and IL-1β, thereby promoting cancer progression^34^. Notably, another ESE subfamily member, ELF3/ESE-1, has been implicated in the regulation of SASP-related genes in senescent cells^43^. These findings suggest that the Ets family proteins, including EHF, participate broadly in senescence-associated inflammatory transcriptional programs, whereas the direction of regulation likely depends on cell type and signaling context.

The clinical relevance of *EHF* expression also appears to be context-dependent. Public cancer resources do not support a common prognostic role for EHF across cancer types; however, individual studies have suggested cancer-type-dependent associations. For example, *EHF* expression has been associated with better prognosis in head and neck squamous cell carcinoma^44,45^. In contrast, high *EHF* expression has been linked to aggressive behavior or poor prognosis in non-small cell lung cancer, colorectal cancer, and thyroid cancer^46–48^. In our GEPIA-based analysis, high *EHF* expression was associated with either favorable, unfavorable, or neutral overall survival depending on tumor type (Supplementary Figure S4), further supporting a context-dependent role of EHF in cancer^49^.

This study also suggests that EHF attenuates paracrine effects that promote migration of neighboring cancer cells (Figure 4). Conditioned medium from *RasG12V*-induced senescent cells promoted migration of HCT116 cells, and this effect tended to be further enhanced when *EHF* was knocked down. In these donor cells, expression of SASP-related genes, including *IL6*, *CXCL8*, and *MMP3,* was increased. However, enhanced cell migration was not observed in conditioned medium from *EHF* knockdown cells when *CXCL8* was not significantly upregulated (Supplementary Figure S3). These observations suggest that cancer cell migration may depend not only on the overall increase in SASP-related genes but also on changes in their composition, potentially including *CXCL8*. Because this study mainly analyzed mRNA expression, the precise contribution of secreted proteins in conditioned medium remains to be elucidated.

Overall, our findings identify EHF as a suppressor of the senescence-associated secretory phenotype in oncogene-induced senescence. Although SASP can confer beneficial effects, such as promoting immune surveillance and tissue repair, its excessive or persistent activation can exacerbate chronic inflammation, age-related diseases, and deterioration of the tumor microenvironment. Therefore, mechanisms that regulate the composition and duration of SASP are likely to be important for controlling the in vivo impact of senescent cells. The EHF-mediated negative regulatory mechanism described here may function as a senomorphic effector that mitigates harmful inflammatory outputs without eliminating senescent cells. Defining how EHF is regulated across cell types and tissue environments may lead to new strategies for controlling persistent inflammation and tumor-promoting SASP driven by senescent cells.

## Acknowledgments

We thank S. Yonehara, T. Kitamura, A. Tugores, E. Nishida, Y. Goto, and M. McMahon for generously providing valuable reagents. We also thank all members of the Ishikawa laboratory at Kyoto University and the Retrotransposon Dynamics laboratory at RIKEN IMS for helpful discussions.

## Funding Information

T.M. was supported by JSPS KAKENHI (23K23863), JST PRESTO (JPMJPR2289), and research grants from the Takeda Science Foundation, the TERUMO Life Science Foundation, the Ichiro Kanehara Foundation for the Promotion of Medical Sciences and Medical Care, the Suzuken Memorial Foundation, the Astellas Foundation for Research on Metabolic Disorders, and the Nagase Science Technology Foundation. F.I. was supported by JSPS KAKENHI (JP19H05655).

## Conflict of Interest

The authors have no conflict of interest.

## Ethics Statement

Approval of the research protocol by an Institutional Review Board: N/A.

Informed Consent: N/A.

Registry and the Registration No. of the study/trial: N/A.

Animal Studies: N/A.

## Author Contributions

Y.F., M.F., F.I., and T.M. conceived and designed the experiments, analyzed data, and prepared the manuscript. Y.F. performed the experiments. Y.K. and Thev. M. provided technical support and contributed to data analysis. Y.F., M.F., F.I., Y.K., Thev. M. and T.M. contributed to critical discussions and to writing and editing the manuscript. All authors approved the final version of the manuscript.

## Data Availability Statement

The data supporting the findings of this study are available in the article and its Supporting Information. Additional raw data are available from the corresponding author upon request.

## List of Supporting Information

Supplementary Figures S1–S4 and Supplementary Tables S1–S2.

---

**Supplementary Figure 1.**
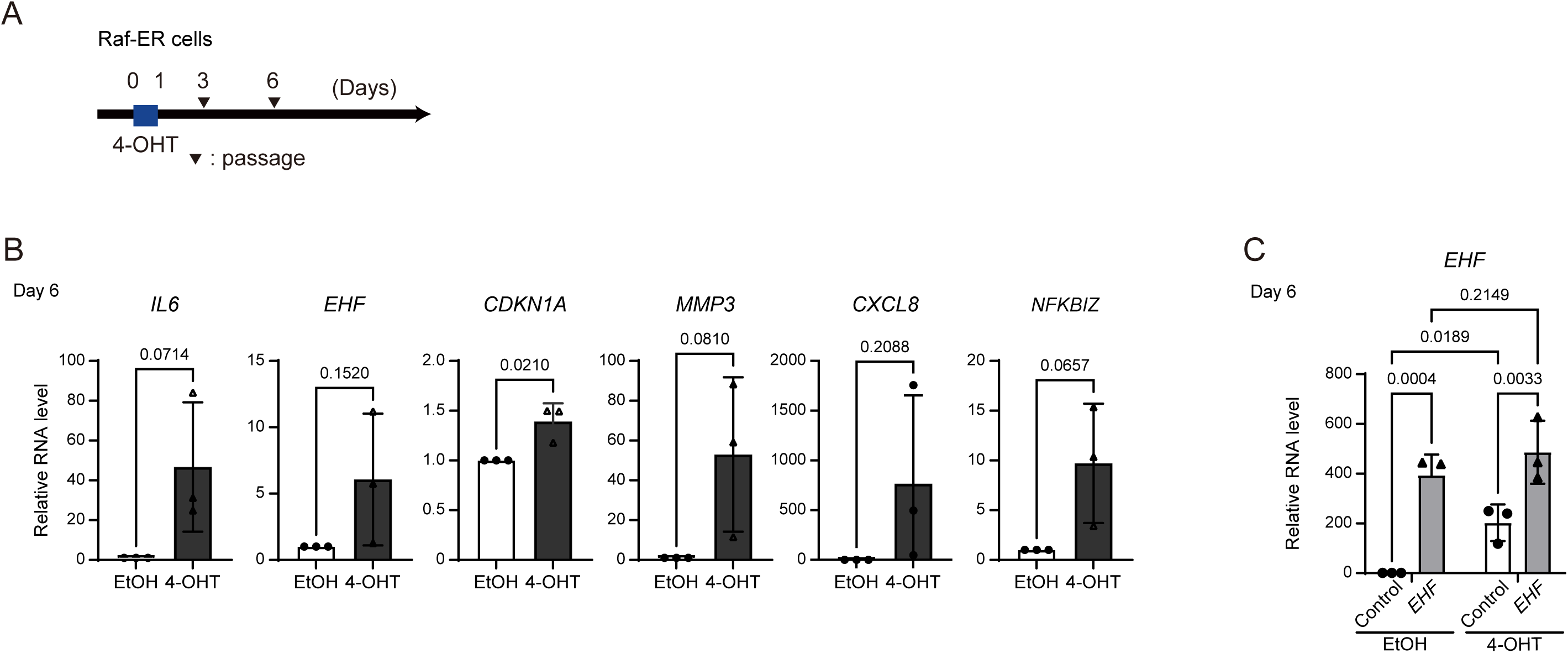

**Supplementary Figure 2.**
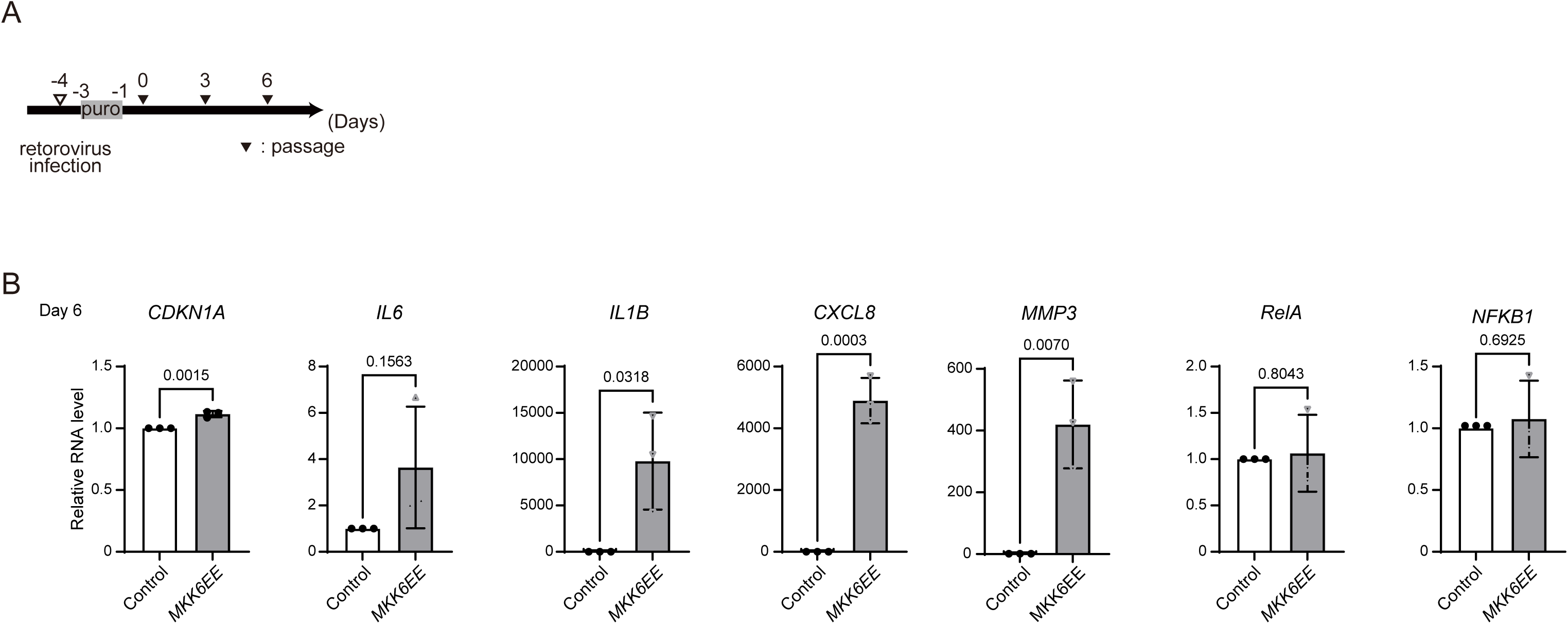

**Supplementary Figure 3.**
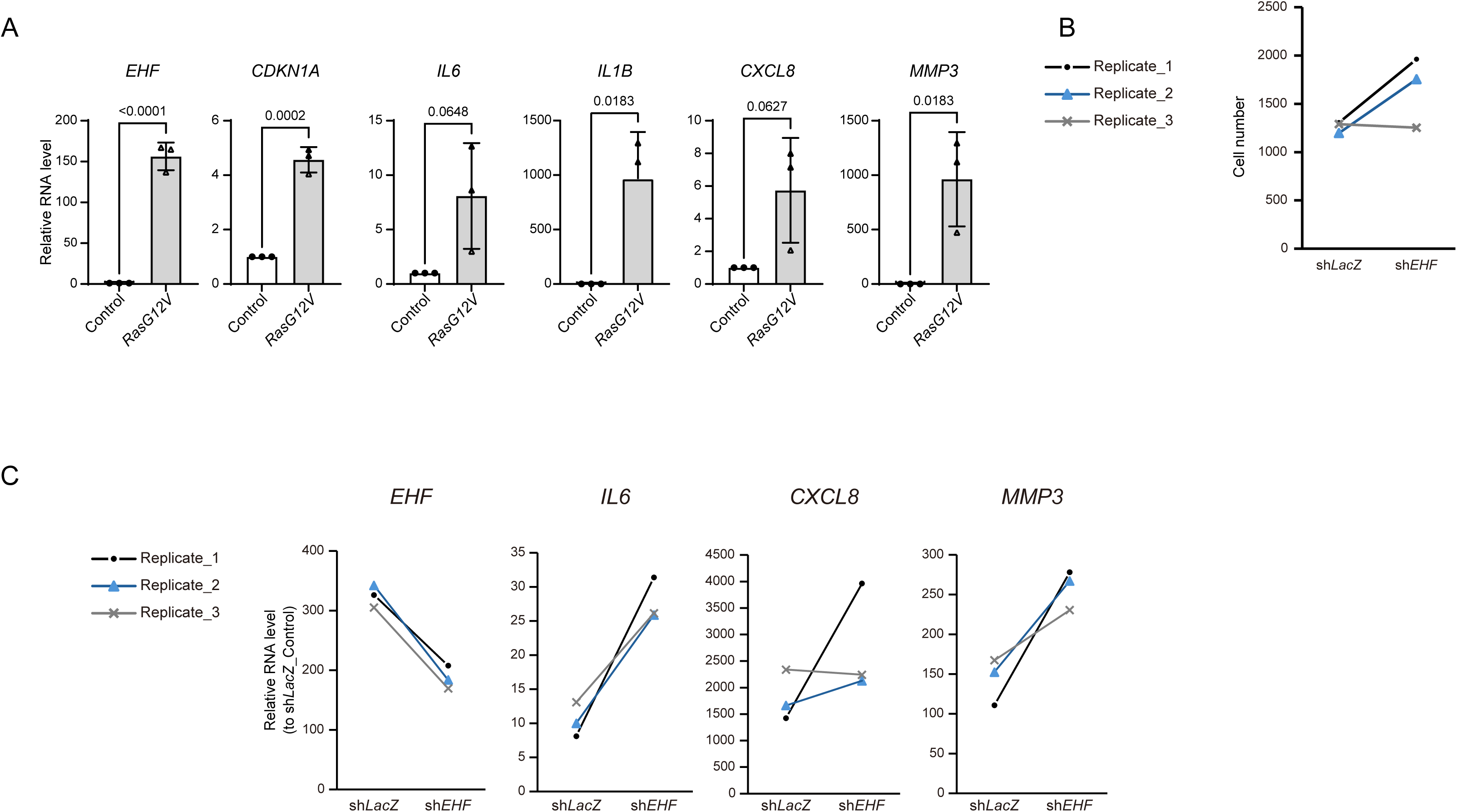

**Supplementary Figure 4.**
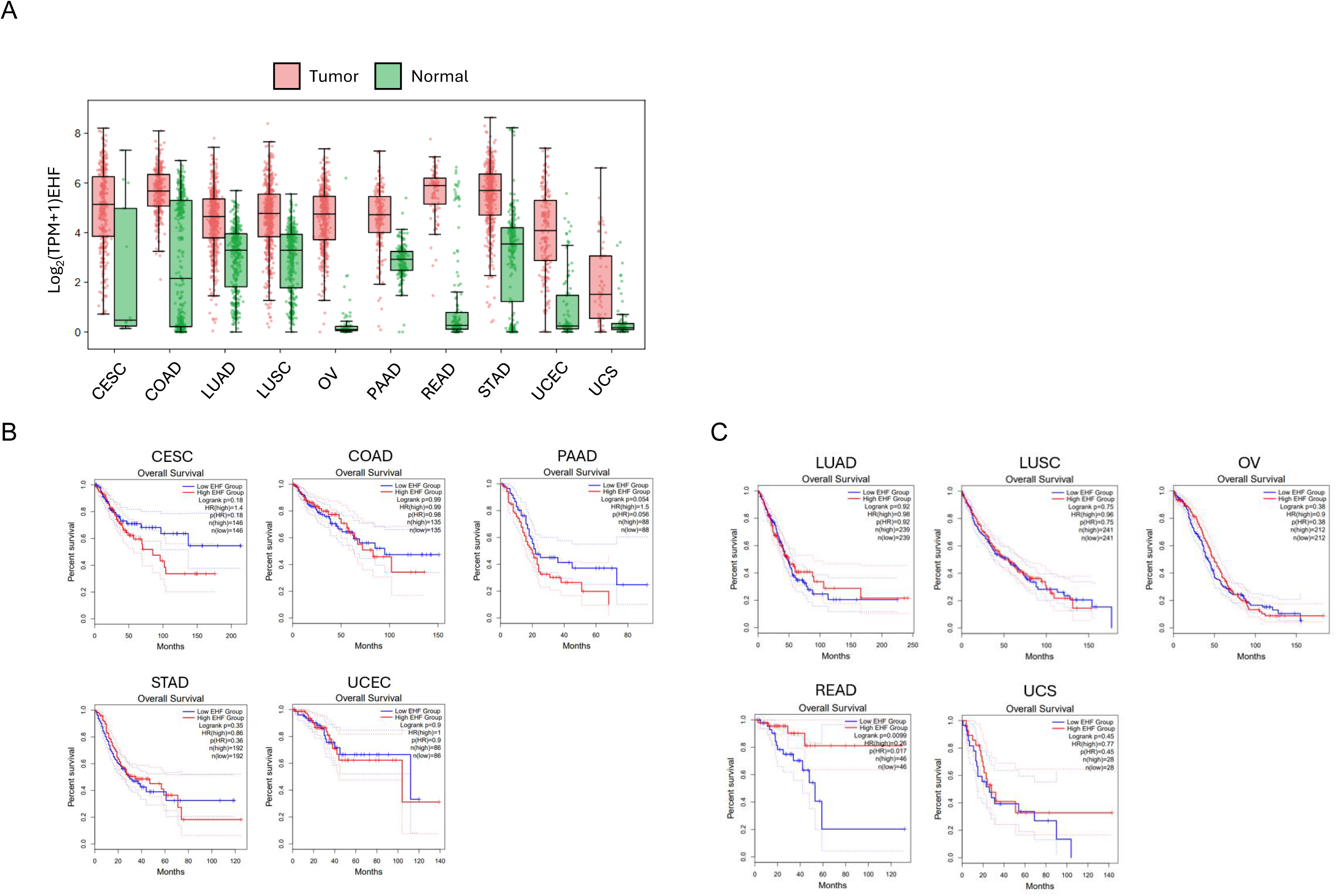

## Notes

### Competing Interest Statement

The authors have declared no competing interest.

