## Supplementary Information for "The Ets family transcription factor EHF suppresses senescence-associated inflammatory responses"

### Contents

Supplementary Figures S1-S4

Supplementary Tables S1-S2

Supplementary References

### Supplementary Figures

**Supplementary Figure S1. Validation of the Raf-ER-inducible senescence system in IMR-90 cells.**

A. Experimental timeline for senescence marker analysis using the Raf-ER system. Cellular senescence was induced in IMR-90 cells expressing the Raf-ER chimeric protein, in which the hormone-binding domain of the estrogen receptor (ER) is fused to the kinase domain of a constitutively activated Raf mutant. Cells were treated with 100 nM 4-hydroxytamoxifen (4-OHT) for 24 hours. As a control, an equal volume of EtOH, the solvent for 4-OHT, was added. The day of 4-OHT or EtOH treatment was designated day 0. Cells were passaged on day 3 and cultured until day 6.

B. RT-qPCR analysis of senescence marker and SASP-related gene expression in the Raf-ER system. RNA was extracted from cells 6 days after Raf-ER-mediated senescence induction, and mRNA levels of SASP-related genes (*IL6*, *CXCL8*, and *MMP3*), *EHF*, *CDKN1A* (p21), and *NFKBIZ* (IκBζ) were measured by RT-qPCR. mRNA levels were normalized to *GAPDH* and shown relative to EtOH-treated cells.

C. RT-qPCR analysis of *EHF* expression in vector-control, WT-EHF-, and ΔETS-EHF-expressing cells after senescence induction using the Raf-ER system. Cells were treated as in A. *EHF* mRNA levels were shown relative to the corresponding EtOH-treated control.

For panels B and C, mRNA levels were normalized to *GAPDH*. Statistical analyses were performed by unpaired t-test for B, and two-way ANOVA followed by Tukey's multiple-comparison test for C. Exact *p*-values are shown in the panel. Error bars indicate SD.

**Supplementary Figure S2. Expression of constitutively active MKK6EE induces senescence- and SASP-related genes**

A. Experimental timeline for MKK6EE-induced cellular senescence in IMR-90 cells. MKK6EE, a constitutively active mutant of MKK6, was introduced into IMR-90 cells via retroviral infection to induce cellular senescence. Cells transduced with the pMX-puro vector served as controls. Cells were collected on day 6 after *MKK6EE* transduction.

B. RT-qPCR analysis of NF-κB- and SASP-related genes after induction of constitutively active MKK6EE in IMR-90 cells. MKK6EE, a constitutively active mutant of MKK6, was introduced into IMR-90 cells by retroviral transduction. Cells transduced with the empty pMX-puro vector served as controls. Cells were collected on day 6 after transduction, and mRNA levels of *CDKN1A*, *IL6*, *IL1B*, *CXCL8*, *MMP3*, *RELA*, and *NFKB1* (p50) were measured by RT-qPCR. mRNA levels were normalized to *GAPDH* and expressed relative to control cells. Statistical analysis was performed by an unpaired t-test. Exact *p*-values are shown in the panel. Error bars indicate SD.

**Supplementary Figure S3. Supplementary data related to the transwell migration assay**

A. RT-qPCR analysis of SASP-related gene expression in IMR-90 donor cells used for the experiment shown in Figure 4B. RNA was isolated from IMR-90 donor cells at the time of conditioned-medium collection, and mRNA levels of *EHF*, *CDKN1A*, and SASP-related genes (*IL6*, *IL1B*, *CXCL8*, and *MMP3*) were measured by RT-qPCR. mRNA levels were normalized to *GAPDH* and shown relative to control cells. Statistical analysis was performed by an unpaired t-test. Exact *p*-values are shown in the panel. Error bars indicate SD.

B. Independent replicate data for the migration assay shown in Figure 4C. Migrated HCT116 cell numbers were compared in each independent experiment (n = 3) using conditioned medium prepared from *RasG12V*-transduced sh*LacZ*- or sh*EHF*-expressing IMR-90 donor cells. The x-axis indicates sample type, and the y-axis indicates migrated cell number.

C. Independent replicate data for the RT-qPCR analysis shown in Figure 4D. mRNA levels of *EHF*, *IL6*, *CXCL8*, and *MMP3* were compared in each independent experiment (n = 3) using IMR-90 donor cells from which conditioned medium was prepared. mRNA levels were normalized to *GAPDH* and shown relative to control-vector-transduced sh*LacZ*-expressing cells. The x-axis indicates sample type, and the y-axis indicates relative mRNA levels.

**Supplementary Figure S4. EHF expression and prognostic significance across epithelial-derived malignancies.**

(A) Differential *EHF* expression between tumor and normal tissues across TCGA cancer cohorts analyzed using GEPIA. *EHF* expression values are shown as log2(TPM + 1). Significant *EHF* overexpression relative to corresponding normal tissues was observed in multiple epithelial-derived malignancies.

(B) Kaplan–Meier overall survival analysis of tumors exhibiting elevated *EHF* expression associated with poor prognosis. High *EHF* expression correlated with significantly reduced survival probability in selected tumor types, indicating context-dependent prognostic roles of EHF across epithelial malignancies.

(C) Kaplan–Meier overall survival analysis of tumors exhibiting elevated *EHF* expression with either favorable or neutral prognostic association. Survival curves were generated using median *EHF* expression as the cutoff value.

**Supplementary Tables**

**Supplementary Table S1. Primer sequences used for RT-qPCR in this study**

| | |
|---|---|
| *CDKN1A* | forward: 5′-AGGTGGACCTGGAGACTCTCAG-3′ |
| | reverse: 5′-TCCTCTTGGAGAAGATCAGCCG-3′ |
| *CXCL8* | forward: 5′-GAGAGTGATTGAGAGTGGACCAC-3′ |
| | reverse: 5′-CACAACCCTCTGCACCCAGTTT-3′ |
| *EHF* | forward: 5′-ATCAGAGGCAGTGGCTCAGCTA-3′ |
| | reverse: 5′-ACCAGTCTTCGTCCATCCACAC-3′ |
| *GAPDH* | forward: 5′-GCAAATTCCTGGCACCCT-3′ |
| | reverse: 5′-TCGCCCCACTTGATTTTGG-3′ |
| *IL1B* | forward: 5′-CCACAGACCTTCCAGGAGAATG-3′ |
| | reverse: 5′-GTGCAGTTCAGTGATCGTACAGG-3′ |
| *IL6* | forward: 5′-AGACAGCCACTCACCTCTTCAG-3′ |
| | reverse: 5′-TTCTGCCAGTGCCTCTTTGCTG-3′ |
| *MMP3* | forward: 5′-GGCAGTTTGCTCAGCCTATC-3′ |
| | reverse: 5′-CAAGGTTCATGCTGGTGTCC-3′ |
| *NFKB1* | forward: 5′-GCAGCACTACTTCTTGACCACC-3′ |
| | reverse: 5′-TCTGCTCCTGAGCATTGACGTC-3′ |
| *NFKBIZ* | forward: 5′-CCGATTCGTTGTCTGATGGACC-3′ |
| | reverse: 5′-GCACTGCTCTCCTGTTTGGGTT-3′ |

*RELA* forward: 5′-TGAACCGAAACTCTGGCAGCTG-3′

reverse: 5′-CATCAGCTTGCGAAAAGGAGCC-3′

oligo(dT) 5′-TTTTTTTTTTTTTTTTTTTTVN-3′

**Supplementary Table S2. Plasmids used in the study**

pMX-puro is a mammalian retroviral expression vector backbone generously provided by Dr. T. Kitamura.

pMX-puro_RasG12V was described previously[1] and contains oncogenic *RasG12V* cloned into pMX-puro.

pMX-puro_MKK6EE was described previously[1–3] and contains constitutively active MKK6 (MKK6EE)-encoding sequence cloned into pMX-puro; *MKK6EE* cDNA was generously provided by E. Nishida and Y. Goto.

pMX-puro_ΔRaf-ER was described previously[2,4] and expresses a chimeric protein consisting of the catalytic domain of constitutively active human Raf-1 (Y340D and Y341D) fused to the hormone-binding domain of the estrogen receptor (ER); ΔRaf-1:ER cDNA was generously provided by Dr. M. McMahon.

pMX-puro_3HA-3FLAG-NFKBIZ contains the *NFKBIZ* coding sequence tagged with 3×HA and 3×FLAG cloned into pMX-puro.

pMXs-neo is a mammalian retroviral expression vector backbone kindly provided by Dr. T. Kitamura.

pMXs-neo_2FLAG-EHF (WT) is a retroviral vector expressing full-length EHF (WT) tagged with 2×FLAG.

pMXs-neo_2FLAG-EHF (ΔETS) is a derivative plasmid of pMXs-neo_2FLAG-EHF (WT), but expresses an EHF mutant lacking the ETS domain (amino acids 1–168) tagged with 2×FLAG.

pCSII-U6-tet-shRNA-neo is a lentiviral vector backbone for shRNA expression generously provided by Dr. S. Yonehara.

pCSII-U6-tet-shRNA-neo_shLacZ is a derivative plasmid of pCSII-U6-tet-shRNA-neo and contains an shRNA sequence targeting *LacZ*.

pCSII-U6-tet-shRNA-neo_shEHF is a derivative plasmid of pCSII-U6-tet-shRNA-neo and contains an shRNA sequence targeting *EHF*.

pCSII-U6-tet-shRNA-neo_shNFKBIZ is a derivative plasmid of pCSII-U6-tet-shRNA-neo and contains an shRNA targeting *NFKBIZ*.

pCMV-VSV-G-RSV-Rev is an expression vector encoding the VSV-G envelope protein and HIV-1 Rev for lentiviral production.

pCAG-HIVgp is an expression vector encoding HIV-1 Gag and Pol for lentiviral packaging.
